# SNED1 fibrillar assembly in the extracellular matrix requires fibronectin and collagen I

**DOI:** 10.64898/2026.03.16.712155

**Authors:** Leanna Leverton, Dharma Pally, Asantewaa Jones, Camille Therol, Sylvie Ricard-Blum, Alexandra Naba

## Abstract

The extracellular matrix (ECM) is a meshwork of proteins that orchestrates a broad range of cellular phenotypes, including proliferation, adhesion, migration, and differentiation. SNED1 is a newly characterized ECM glycoprotein that promotes cell adhesion and is essential for embryonic development. Its upregulation is also associated with breast cancer metastasis and poor prognosis for breast cancer patients. We recently showed that SNED1 assembles into fibrillar structures, but the mechanisms guiding its incorporation into the ECM scaffold remain unknown. Combining biochemical assays and confocal immunofluorescence imaging, we found that SNED1 assembly in the ECM occurs early in the process of ECM building and is concomitant and overlaps with the deposition of fibronectin and collagen I, two major ECM proteins. By knocking down fibronectin or destabilizing collagen I fibers, we further demonstrate that SNED1 requires the presence of these proteins for its assembly. Last, using biolayer interferometry, we identify collagen I as the first direct binding partner of SNED1. Altogether, our results lay the foundation for future studies aimed at determining the mechanisms by which SNED1 fibers contribute to SNED1 pathophysiological functions.

**SUMMARY STATEMENT:** The novel protein SNED1 requires the presence of fibronectin and collagen I to assemble into fibrillar structures in the extracellular matrix scaffold.

## INTRODUCTION

The extracellular matrix (ECM) is a complex and tightly regulated assembly of over 150 proteins that provides physical support to surrounding cells (Karamanos et al., 2021; Naba, 2024). The ECM also exerts signaling functions that control a broad range of cellular processes, including proliferation (Hastings et al., 2019), adhesion, and migration (Pally and Naba, 2024; Yamada and Sixt, 2019). Alterations of the structural integrity of the ECM have been associated with a plethora of clinical manifestations spanning hereditary diseases (Lamandé and Bateman, 2020), impaired tissue repair (Barker and Engler, 2017; Midwood et al., 2004), cancer (Cox, 2021), and fibrosis (Ricard-Blum et al., 2018). The process of ECM assembly relies extensively on biomolecular interactions, such as those between ECM proteins and their cell-surface receptors, as extensively described for one of the most abundant ECM proteins, fibronectin, and its receptors of the integrin family (Mao and Schwarzbauer, 2005; Sun et al., 2025). ECM build-up also relies on interactions between ECM proteins. For example, fibronectin has been shown to act as a scaffolding protein allowing the assembly of collagen I or fibrillins, themselves facilitating the incorporation of additional proteins like other fibrillar and non-fibrillar collagens or latent TGFβ-binding proteins (Canty and Kadler, 2005; Handford et al., 2000; Kadler et al., 2008; Sabatier et al., 2009; Schwarzbauer and DeSimone, 2011; Singh et al., 2010; Sun et al., 2025). Gaining insights into the process of ECM fibrillogenesis and the interactions involved in ECM assembly is thus key to understanding the multiple roles of ECM proteins in health and disease.

We have recently characterized a new ECM protein called SNED1 (Sushi, Nidogen, and EGF-like Domains). SNED1 promotes cell adhesion (Pally et al., 2025) and is essential for embryonic development, as knocking out *Sned1* in mice resulted in early neonatal lethality, in part due to craniofacial malformations (Barqué et al., 2021). SNED1 upregulation is also associated with breast cancer metastasis and poor prognosis for breast cancer patients (Naba et al., 2014). We also recently observed that SNED1 assembles into fibrillar structures in the ECM (Vallet et al., 2021) and have predicted that SNED1 could interact with a broad range of ECM glycoproteins, including fibronectin and collagens (Vallet et al., 2021), although, to date, no direct ECM interactor of extracellular SNED1 has been reported experimentally. We also do not know yet the mechanisms guiding SNED1’s incorporation into the ECM scaffold.

In this short report, we describe the establishment of a cell culture system employing *Sned1^KO^* immortalized embryonic fibroblasts overexpressing SNED1-GFP to generate ECMs in which we can visualize SNED1 incorporation *in vitro*. Combining biochemical assays and confocal immunofluorescence imaging, we found that SNED1 assembly in the ECM occurs early in the process of ECM formation and is concomitant to and overlaps with the deposition of fibronectin and collagen I. By knocking down fibronectin using short interfering RNA or altering the collagen I meshwork, we demonstrate that SNED1 requires the presence of these two proteins for its assembly and incorporation into the ECM. Last, using biolayer interferometry, we identify collagen I as the first direct binding partner of SNED1. Altogether, our results lay the foundation for future studies aimed at determining the mechanisms by which SNED1 fibers contribute to SNED1’s functions in development and metastasis.

## RESULTS AND DISCUSSION

### SNED1 assembles in the ECM over time and adopts a fibrillar pattern

To study the mechanisms leading to SNED1 assembly in the ECM, we established an *in-vitro* system using *Sned1^KO^* immortalized mouse embryonic fibroblasts (hereafter referred to as *Sned1^KO^* iMEFs) stably overexpressing SNED1-GFP (**Figure 1A**). Using immunoblotting, we first confirmed that these cells expressed SNED1 (**Figure 1B, left panel**) and secreted the protein by monitoring its presence in the culture medium (**Figure 1B, right panel**). We next confirmed that SNED1 could assemble into the ECM. To do so, we performed a deoxycholate (DOC) solubility assay and found that, in addition to being present in the DOC-soluble fraction, representative of intracellular proteins (**Figure 1C, left panel**), SNED1 was also present in the DOC-insoluble fraction, enriched for ECM proteins, such as collagen I, as early as 3 days post-seeding (**Figure 1C, right panels**). Interestingly, we consistently observed a second, higher molecular weight band for SNED1 in the DOC-insoluble fraction when cells are in culture for a longer period (> 6 days post-seeding). This result suggests that different proteoforms of SNED1, possibly resulting from post-translational modifications, arise extracellularly as SNED1 is incorporated in the insoluble ECM meshwork. It would be interesting in the future to clarify the biochemical nature of these proteoforms and their significance.

**Figure 1.**
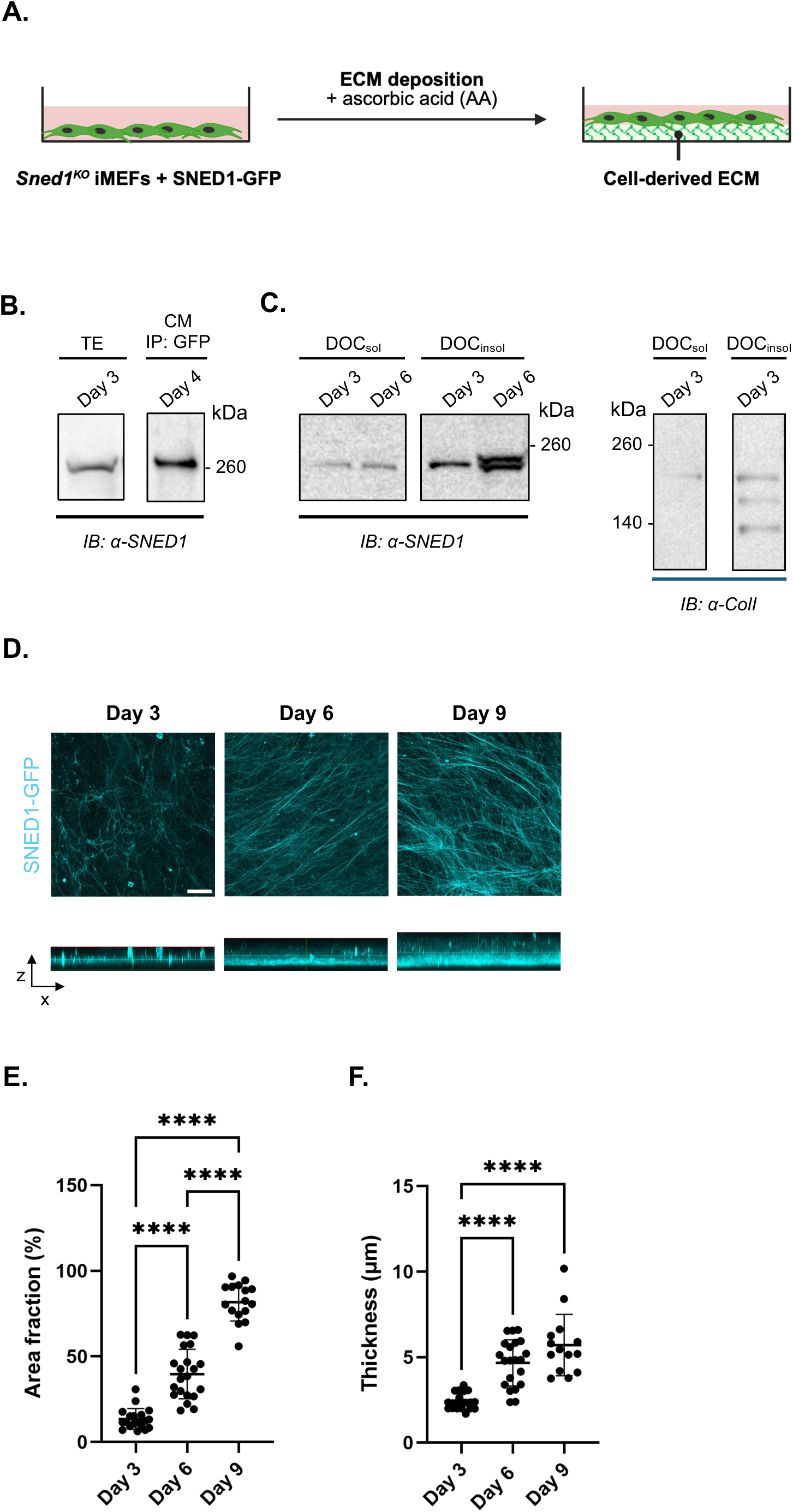
SNED1 assembles in the insoluble ECM over time. **A.** Schematic depiction of the workflow used to generate cell-derived ECMs. *Sned1^KO^*iMEFs overexpressing SNED1-GFP are seeded on glass coverslips coated with gelatin and stimulated with ascorbic acid (AA) as they deposit ECM over time. *Created in BioRender. Leverton, L. (2026)* https://BioRender.com/xnqn90a. **B.** Immunoblots using an anti-SNED antibody show SNED1-GFP in total protein extracts (TE) and culture medium (CM) conditioned by cells cultured for 3 or 4 days post-seeding. Images are representative of at least two biological replicates. **C. *Left panels***: Immunoblots using an anti-SNED1 antibody show SNED1-GFP in the DOC-soluble (DOC_sol_) and DOC-insoluble (DOC_insol_) protein fractions collected from *Sned1^KO^*iMEFs overexpressing SNED1-GFP 3 or 6 days post-seeding. ***Right panels***: Immunoblots show collagen I in the DOC-soluble (DOC_sol_) and DOC-insoluble (DOC_insol_) protein fractions collected from *Sned1^KO^* iMEFs overexpressing SNED1-GFP 3 days post-seeding. The different bands correspond to different proteoforms and chains (α1 and α2) of collagen I. Images are representative of at least two biological replicates. **D.** Orthogonal maximum projections (XY, *top panels*; XZ, *bottom panels*) show SNED1 fibers in decellularized ECMs produced by cells 3, 6, or 9 days post-seeding. Images are representative of three independent biological replicates. Scale bar: 20 µm. **E.** Dot plot shows the percentage of SNED1 signal area fraction at each timepoint. Individual experimental values from three independent biological replicates, with at least 4 imaging fields per replicate and timepoint, are represented together with the mean value (black bar) ± standard deviation. ****p<0.0001 using Welch and Brown-Forsythe one-way ANOVA with Dunnett’s T3 correction for multiple comparisons. **F.** Dot plot shows the thickness of the SNED1 signal (µm) at each timepoint. Individual experimental values from three independent biological replicates, with at least 4 imaging fields per replicate and timepoint, are represented together with the mean value (black bar) ± standard deviation. ****p<0.0001 using Welch and Brown-Forsythe one-way ANOVA with Dunnett’s T3 correction for multiple comparisons.

Additionally, we confirmed by immunoblotting that the overexpression of SNED1 did not alter the abundance of the two most abundant fibrillar ECM proteins, fibronectin and collagen I (**Supplemental Figure S1A**). Using confocal imaging microscopy, we also established that *Sned1^KO^* iMEFs overexpressing SNED1-GFP produced an ECM overall similar in density and thickness than the ECMs produced by wild-type iMEFs or by *Sned1^KO^* iMEFs overexpressing GFP alone (**Supplemental Figures S1B and S1C**), although the fibrillar meshwork of ECM proteins produced by wild-type iMEFs seemed to be somewhat slightly less aligned (**Supplemental Figure S1D**). We also observed that the colocalization of fibronectin and collagen I was unchanged across conditions (**Supplemental Figure S1B**).

With this experimental system in place, we first sought to characterize the incorporation and pattern of distribution of SNED1 in the ECM. For that, we decellularized cell-derived ECMs at different culture time points, with day 3 representing the early step of ECM assembly and days 6 and 9 selected to capture ECM maturation and remodeling. Imaging of the decellularized ECMs revealed the presence of a thin layer of SNED1-GFP fibers as early as 3 days post-seeding (**Figure 1D, left panels**). The density of SNED1 fibers, determined through the calculation of area fraction, increased from day 3 to day 6 post-seeding (**Figure 1D, middle panels**) and further increased by day 9 post-seeding (**Figure 1D, right panels**), as quantified in **Figure 1E**. The thickness of the SNED1 layer within the ECM also increased over time, more significantly between day 3 and day 6, and with a trend toward increase between day 6 and day 9 (**Figures 1D and 1F).** Altogether, these data show that SNED1 is incorporated into the insoluble ECM meshwork as fibrillar structures, and the density of SNED1-GFP staining increases over time.

### SNED1 colocalizes with fibronectin and collagen I at the early stage of ECM assembly, but not in mature ECMs

One of the main mechanisms leading to protein fibrillar assembly into the ECM is via ECM protein-ECM protein interactions, as extensively described for fibronectin, collagens, and elastic fibers (Godwin et al., 2019; Handford et al., 2000; Holmes et al., 2018; Naba, 2024; Singh et al., 2010; Singh et al., 2021; Sun et al., 2025). We previously identified fibronectin, a protein known for its scaffolding ability (Singh et al., 2010), as a putative binding partner of SNED1 (Vallet et al., 2021). We thus sought to determine the extent of the colocalization between SNED1 and fibronectin over the course of ECM assembly and maturation.

Using confocal fluorescence microscopy, we observed that SNED1 and fibronectin staining colocalized in decellularized ECMs produced by cells as early as 3 days post-seeding (**Figure 2A, left panels and Figure 2B, left panel**). We also observed that the relative abundance of SNED1 and fibronectin fibrils (**Figure 2A**) and the thickness of the layers formed by these respective meshworks increased over time (**Figures 2C and 2D**). However, surprisingly, rather than increasing, the colocalization between SNED1 and fibronectin decreased over time, as determined by the calculation of Manders correlation coefficient (**Figure 2B, middle and right panels**), to the point of forming gradually two distinct layers within the ECM, as visualized on XZ projections of confocal imaging stacks (**Figure 2C, right panel**). We sought to further quantify this phenotype by plotting the means of the lowest and of the highest Z-slices within the imaging stacks showing in-focus staining for SNED1 and fibronectin, respectively, normalized all values to SNED1’s lowest Z-slice value, and estimated the thickness of the layers in which SNED1 and fibronectin overlapped (**Figure 2D**). At day 3, the overlap between SNED1 and fibronectin spanned the full thickness of the ECM (**Figure 2D, left panel**). However, by day 6, we observed that the upper mean for fibronectin began shifting upward (**Figure 2D, middle panel**). At day 9, we observed a stratification of the ECM between a fibronectin-rich layer and a SNED1-rich layer, with the most basal portion of the ECM (∼1.75 µm) being statistically significantly enriched for SNED1 and depleted for fibronectin (**Figure 2D, right panel**), while the most apical portion of the ECM meshwork (∼1.5 µm) tended to contain fibronectin and devoid of SNED1, although this was not statistically significant. Importantly, this partitioning did not result from multiple cell layers, as staining of the cellular culture (cells and their ECM) revealed that the cells, even after 9 days in culture, remained largely organized into a monolayer (**Supplemental Figure S2A**).

**Figure 2.**
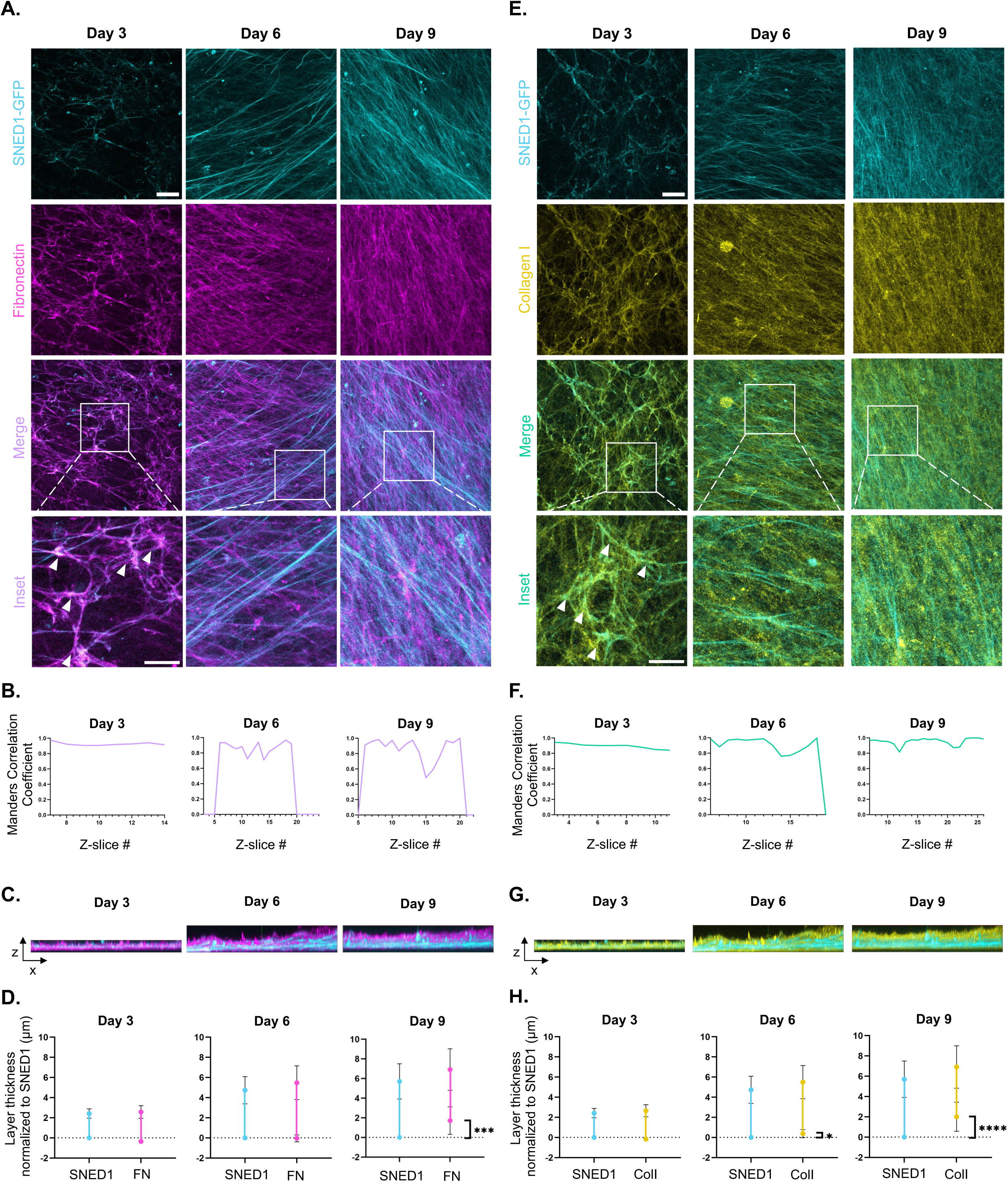
SNED1 colocalizes with fibronectin and collagen I during ECM assembly. **A, E.** XY orthogonal maximum projections show SNED1 (cyan) and fibronectin (FN; magenta), or collagen I (yellow) fibers in the ECM produced by cells and decellularized 3, 6, or 9 days post-seeding. Merge panels and insets show overlap (white arrow heads) between SNED1 and fibronectin (**A**) or collagen I (**E**). Scale bar: 20 µm and inset scale bar: 10 µm. Images are representative of at least three independent biological replicates. **B.** Line graphs show, for each timepoint, the Manders correlation coefficient between SNED1 and fibronectin signals for each Z-slice of a representative imaging field. **C.** XZ projections show the organization of SNED1 and fibronectin layers within the thickness of decellularized ECMs at each timepoint. Images are representative of at least three independent biological replicates. **D.** Graphs depict the overlap of the SNED1 and fibronectin (FN) signals (µm) at each timepoint in the XZ dimension. Lower data point represents the mean value (± s. d.) of the position of the lowest Z planes where in-focus signal was detected; the upper data point represents the mean value (±s. d.) of the position of the highest Z planes where in-focus signal was detected. Experimental values from three biological replicates, with at least 4 imaging fields per biological replicate and timepoint, were included in the calculation. Values were normalized to SNED1’s lowest Z-slice value. ***p<0.001 using unpaired t-test with Welch’s correction. **F.** Line graphs show, for each timepoint, the Manders overlap coefficient between the SNED1 and collagen I signals for each Z-slice of a representative imaging field. **G.** XZ projections show the spatial organization of SNED1 and collagen I layers within the thickness of decellularized ECMs at each timepoint. Images are representative of at least three independent biological replicates. **H.** Graphs depict the overlap of SNED1 and collagen I (ColI) signals (µm) at each timepoint in the XZ dimension. Lower data point represents the mean value (± s. d.) of the position of the lowest Z planes where in-focus signal was detected; the upper data point represents the mean value (±s. d) of the position of the highest Z planes where in-focus signal was detected. Experimental values from three biological replicates, with at least 4 imaging fields per biological replicate and timepoint, were included in the calculation. Values were normalized to SNED1’s lowest Z-slice value. *p<0.05, ****p<0.0001 using unpaired t-test with Welch’s correction.

Fibronectin is a master organizer of ECM architecture, and its protein-domain-based organization constitutes a scaffolding platform to support the assembly of other ECM proteins (Naba, 2024; Singh et al., 2010), including the most abundant protein of the ECM, the fibrillar collagen, collagen I (Kadler et al., 2008; Paten et al., 2019; Saunders and Schwarzbauer, 2019; Sottile et al., 2007). We thus next sought to determine the extent of the colocalization between SNED1 and collagen I. Similar to our observations with fibronectin, we found that SNED1 and collagen I staining overlapped in newly assembled ECMs (**Figures 2E and 2F**). As expected, the density of collagen I fibers (**Figures 2E and 2F**) and the thickness of the collagen I meshwork (**Figures 2G and 2H**) increased over time. However, and as observed with fibronectin, as ECMs matured, the SNED1 fiber meshwork began to partition from the collagen I meshwork at day 6, forming distinct SNED1-positive and collagen-I-positive layers in the ECM at day 9 (**Figures 2G and 2H**). Specifically, we observed a shift in both the lower and upper mean focal planes positive for collagen I at day 6, with statistically different lower mean positions between SNED1 and collagen I (**Figure 2H, middle panel**). At day 9, the quantification of the stratification observed in the XZ projections (**Figure 2G, right panel**) revealed that the most basal portion of the ECM (∼lower 2 µm) contained only SNED1 signal (**Figure 2H**) and that the collagen-I-positive layer started at a statistically significant higher level within the ECM (**Figure 2H**). Importantly, we confirmed that fibronectin and collagen I remained tightly associated throughout ECM assembly and maturation in our experimental system (**Supplemental Figures S2B and S2C**).

Taken together, these results show an association of SNED1 with the two most abundant fibrillar proteins of the ECM, fibronectin and collagen I, at early stages of ECM assembly. However, this association is only transient and, as ECMs mature, a partitioning of the SNED1 and fibronectin/collagen I meshworks occurs. Understanding the mechanistic bases of this latter observation is the focus of the companion paper by Pally, Leverton, *et al*., submitted in conjunction with this paper.

### Fibronectin knockdown disrupts SNED1 ECM assembly

Since we observed a colocalization between SNED1 and fibronectin at the early step of ECM assembly, we next asked whether SNED1 assembly requires the presence of fibronectin. To test this, we used an RNA interference strategy to knock down fibronectin (*Fn1*) expression in *Sned1^KO^* iMEFs overexpressing SNED1-GFP. Since fibronectin is present in the serum, and cells can pull down fibronectin molecules from the culture medium via integrin receptors to initiate fibrillogenesis, we cultured the cells with a medium containing fibronectin-depleted serum. In addition, to further limit fibronectin assembly, the cells were seeded on Matrigel, a basement-membrane-type of ECM devoid of collagen I and fibronectin rather than on gelatin.

Staining of the ECMs produced by cells for 3 days confirmed a statistically significant decrease in the fibronectin signal upon *Fn1* knockdown, as compared to the staining observed in ECM produced by cells transfected with a control siRNA (**Figures 3A and 3B**). We also observed that si#1 and si#2 led to a larger decrease in the number of fibronectin fibers formed than si#3 (**Figures 3A**). Importantly, we observed a concomitant, and statistically significant, reduction in the number and density of SNED1 fibrils (**Figures 3A and 3B**). This observation suggests that SNED1 requires fibronectin for its fibrillar assembly in the ECM.

**Figure 3.**
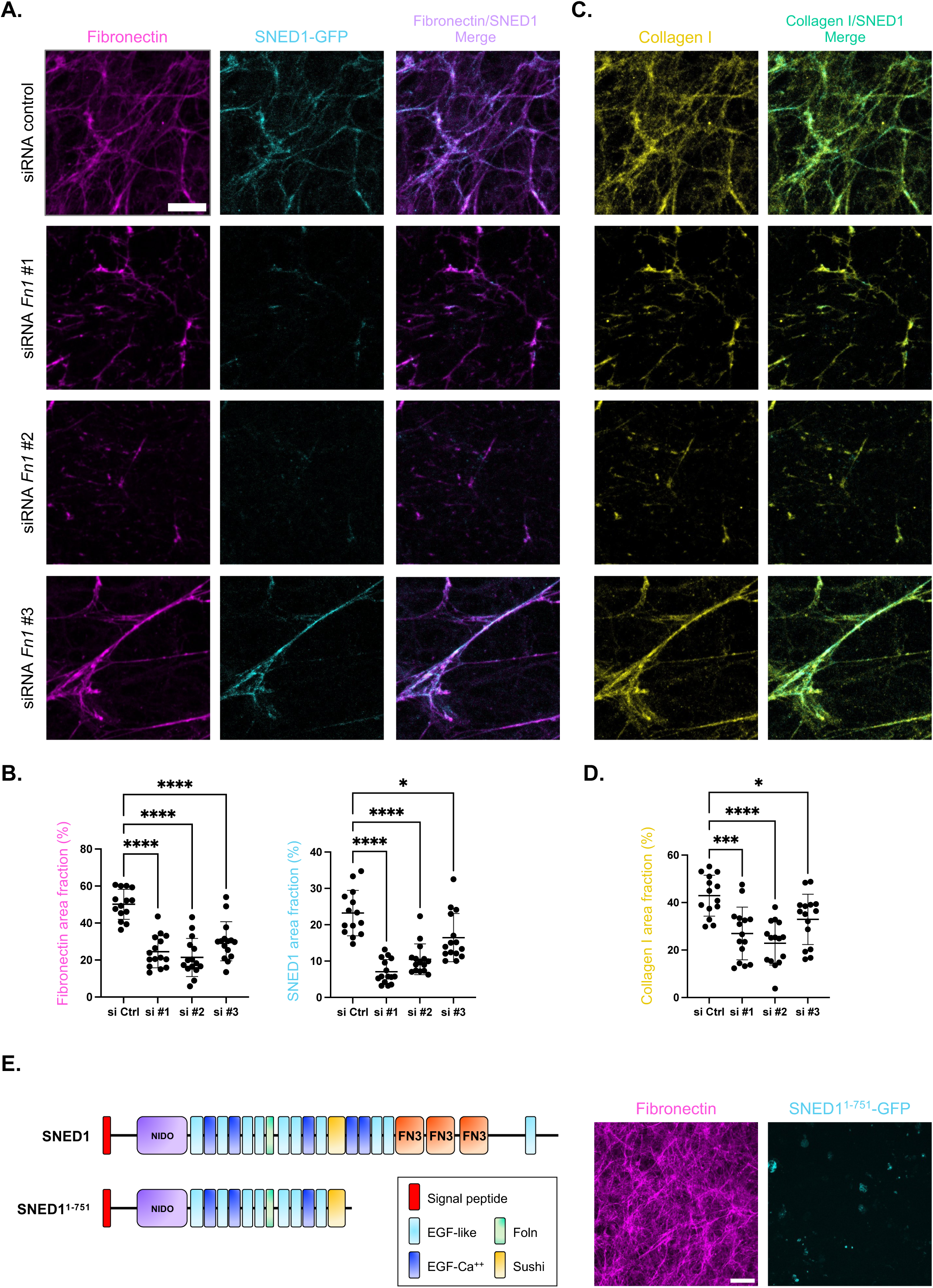
Fibronectin knockdown disrupts SNED1 fibrillar assembly. **A.** XY orthogonal maximum projections show fibronectin (magenta), SNED1 (cyan), and the overlap between the fibronectin and SNED1 signals (merge; purple) in the ECM produced by *Sned1^KO^* iMEFs overexpressing SNED1-GFP and decellularized 3 days post-transfection. Conditions include cells transfected with siRNA control (*top panels*) or independently with three different siRNA constructs targeting fibronectin (si#1, si #2, or si #3; *lower three panels*). Images are representative of three biological replicates. Scale bar: 10 µm. **B.** Dot plot shows the percentage of fibronectin (*left panel*) or SNED1 (*right panel*) signal area fraction for each condition. Individual experimental values from three independent biological replicates, with at least 4 imaging fields per replicate, are represented together with the mean value (black bar) ± standard deviation. *p<0.05, ****p<0.0001 using Welch and Brown-Forsythe one-way ANOVA with Dunnett’s T3 correction for multiple comparisons. **C.** XY orthogonal maximum projections show collagen I (yellow) and overlap between the fibronectin (*see panels in **A***) and collagen I signals (merge; green) in the ECM produced by *Sned1^KO^* iMEFs overexpressing SNED1-GFP and decellularized 3 days post-seeding. Conditions include cells transfected with a scrambled siRNA control (*top panels*) or independently with three different siRNA constructs targeting fibronectin (*lower three panels*). Images are representative of three biological replicates. Scale bar: 10 µm. **D.** Dot plot shows the percentage of collagen I signal area fraction for each condition. Individual experimental values from three independent biological replicates, with at least 4 imaging fields per replicate, are represented together with the mean value (black bar) ± standard deviation. *p<0.05, ***p<0.001****p<0.000 using Welch and Brown-Forsythe one-way ANOVA with Dunnett’s T3 correction for multiple comparison). **E. *Left panel:*** Schematic representation of the domain organization of human SNED1 and its truncated form (SNED1^1-751^) lacking the C-terminal fibronectin type III (FN3) domains (orange); ***Right panel:*** XY orthogonal maximum projections show fibronectin (magenta) and SNED1 (cyan) in the ECM produced by *Sned1^KO^* iMEFs overexpressing SNED1^1-751^-GFP and decellularized 9 days post-seeding. Scale bar: 20 µm.

However, we also observed a decrease in the density of collagen I fibers upon *Fn1* knockdown (**Figures 3C and 3D**). This result is not surprising, since it is well established that collagen I fibrillogenesis requires fibronectin (Kadler et al., 2008; Saunders and Schwarzbauer, 2019; Sottile et al., 2007); however, it raises the possibility that, instead of depending directly on the presence of fibronectin, SNED1 fibrillar assembly into the ECM is primarily dependent on collagen I. One way to address this is to determine whether SNED1 can interact with fibronectin, as our *in-silico* predictions suggested (Vallet et al., 2021). However, neither co-immunoprecipitation performed on the conditioned medium of cells overexpressing SNED1-GFP nor biolayer interferometry (BLI) yielded conclusive results regarding SNED1’s ability to bind fibronectin (*data not shown*). Importantly, the techniques we used assess the ability of purified soluble proteins to interact *in vitro*; as such, the negative results we obtained do not fully rule out that these proteins could interact within the ECM, where proteins adopt conformations that differ from that of those of their soluble forms (Leverton et al., 2026).

Since SNED1 and fibronectin were predicted to interact through fibronectin type III (FN3) domains (Vallet et al., 2021), we sought to determine whether the FN3 domains in SNED1 were required to mediate SNED1 assembly into the ECM. Interestingly, we found that deletion of the C-terminal region of SNED1, which contains the three FN3 domains, abolished the fibrillar incorporation of SNED1 in the ECM (**Figure 3E**).

### SNED1 fibrillar assembly in the ECM requires collagen I

Since *Fn1* knockdown altered the assembly of the collagen I meshwork, we further wanted to delineate whether SNED1 requires collagen I in the ECM for its assembly. Ascorbic acid (vitamin C) is a cofactor for lysyl hydroxylases and prolyl hydroxylases, two families of enzymes playing key roles in post-translational modifications of collagens and in the stabilization of the collagen triple helix. Evidence also shows that ascorbic acid can modulate several aspects of collagen I biosynthesis, including gene expression, hydroxylation, folding, secretion, and cross-linking by modulating the activity of lysyl oxidase (Gjaltema and Bank, 2017; Murad et al., 1981; Pinnell, 1985; Rappu et al., 2019; Salo and Myllyharju, 2021; Schwarz, 1985; Schwarz, 2015). We thus omitted ascorbic acid from our cell culture system to reduce collagen I.

Immunoblot analyses on total protein extracts from cells treated or not with ascorbic acid revealed that the absence of ascorbic acid from the culture medium led to an increase in certain proteoforms of collagens (in particular those, based on molecular weight corresponding to intracellular immature pro-collagens, that are likely be under-hydroxylated and thus destined to degradation in line with published observations (Canty and Kadler, 2005)). Conversely, we observed an increase in high-molecular-weight collagen polymers in total protein extracts that likely correspond to the pool of ECM-assembled collagen I (**Figure 4A, left panel**). We also detected a slightly elevated amount of mature collagen I in the conditioned cell culture medium in the presence of ascorbic acid (**Figure 4A, middle panel; red asterisk**) and observed a large increase in the number of high molecular-weight polymers in the DOC-insoluble protein fraction of cells grown in the presence of ascorbic acid (**Figure 4A, right panels; red asterisk**). These observations suggest that in our experimental system, the absence of ascorbic acid, indeed alters collagen I meshwork. We further confirmed these observations using confocal immunofluorescence microscopy where the density of collagen I fibers deposited in the ECM was statistically significantly decreased in absence of ascorbic acid (**Figures 4B and 4C, left panels**). Importantly, we found that the alteration of the collagen I meshwork was accompanied by a concomitant, and statistically significant, decrease in SNED1 staining density (**Figures 4B and 4C, middle panels**). Since the absence of ascorbic acid did not lead to a complete absence of collagen deposition, we also observed a steady association of SNED1-positive fibers with the remaining collagen I fibrils.

**Figure 4.**
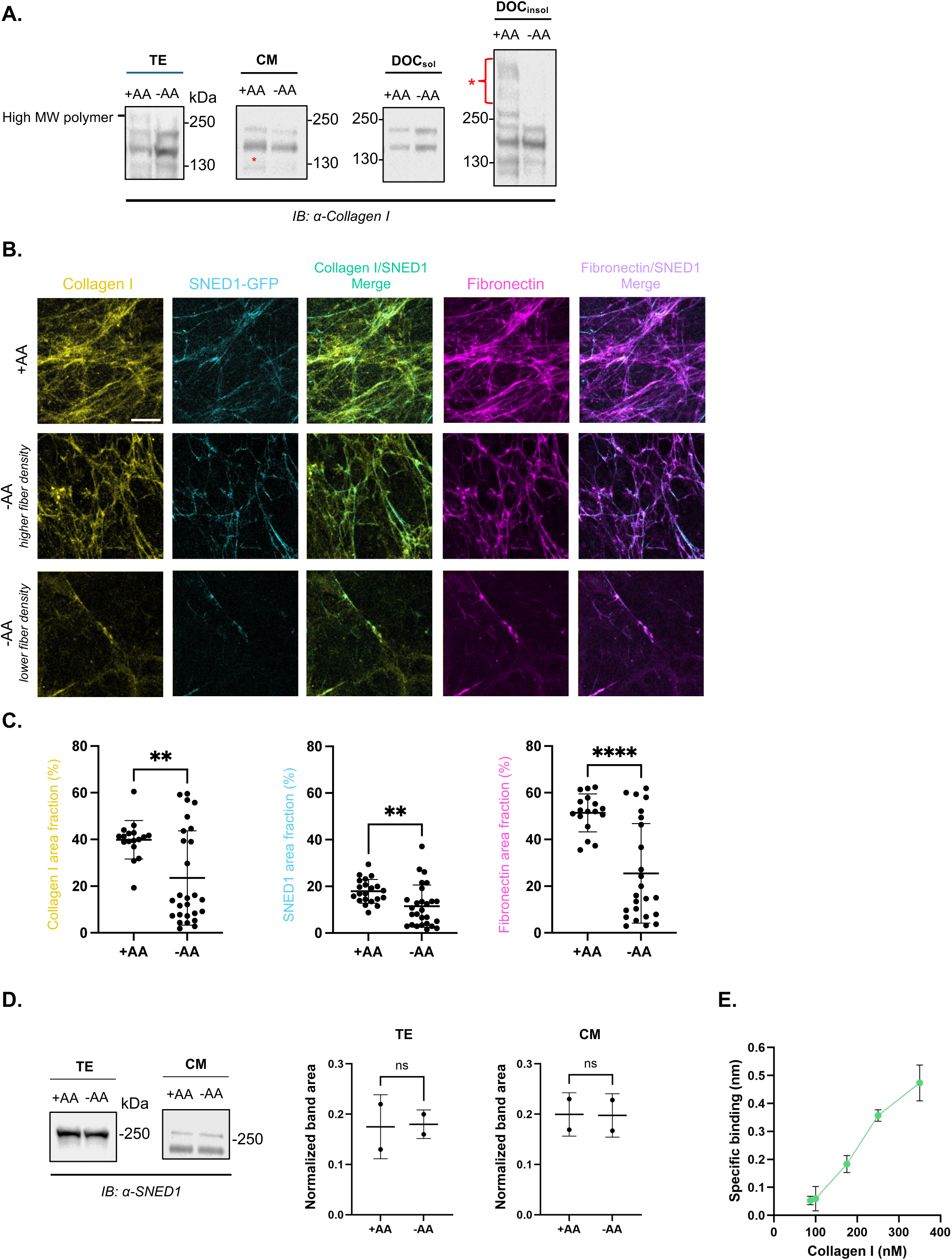
SNED1 assembly requires the presence of collagen I. **A.** Immunoblots show the presence of collagen I in total protein extracts (TE), conditioned medium (CM), and DOC-soluble (DOC_sol_) and DOC-insoluble (DOC_insol_) protein fractions collected from *Sned1^KO^*iMEFs overexpressing SNED1-GFP cultured either in the presence (+AA) or absence (-AA) of ascorbic acid. Red asterisks indicate the increased abundance of higher molecular weight assembled collagen polymers in the presence of ascorbic acid. Images are representative of at least two biological replicates. **B.** XY orthogonal maximum projections show collagen (yellow), SNED1 fibers (cyan), and fibronectin (magenta) in the ECM produced by *Sned1^KO^*iMEFs overexpressing SNED1-GFP treated with (+AA) or without (-AA) ascorbic acid (with representative images of fields presenting higher or lower collagen I fiber density in the -AA condition), 3 days post-seeding. Images are representative of three biological replicates. Scale bar: 10 µm. **C.** Dot plot shows the percentage of collagen I (*left panel*), SNED1 (*middle panel*), and fibronectin (*right panel*) signal area fractions, for each condition. Individual experimental values from three independent biological replicates, with at least 4 imaging fields per replicate, are represented together with the mean value (black bar) ± standard deviation. **p<0.01, ****p<0.0001 using unpaired t-test with Welch’s correction. **D. *Left panels:*** Immunoblots using an anti-SNED1 antibody show the presence of SNED1-GFP in total protein extracts (TE), conditioned medium (CM), and DOC-soluble (DOC_sol_) and DOC-insoluble (DOC_insol_) protein fractions collected from *Sned1^KO^* iMEFs overexpressing SNED1-GFP cultured either in the presence (+AA) or absence (-AA) of ascorbic acid. Images are representative of two biological replicates. ***Right panels:*** Bar charts represent the quantification of immunoblot signal intensity, where the area of the band corresponding to SNED1 in total protein extracts or the conditioned culture media of cells cultured either in the presence (+AA) or absence (-AA) of ascorbic acid was normalized to total protein content determined by Ponceau staining. Unpaired t-test with Welch’s correction did not show statistical significance (ns). **E.** Line graph depicts specific binding identified by biolayer interferometry of acid-soluble human collagen I to immobilized Sned1, at increasing concentration of collagen I. Data show mean ± standard deviation of three independent experimental replicates.

However, our experiments also show that altering the collagen I meshwork leads to a significant decrease in fibronectin fibrils (**Figures 4B and 4C, right panels**), further highlighting the tight interplay between these two proteins (*see above*). Of note, the abundance of SNED1 produced and secreted by cells was unaffected by absence of ascorbic acid, confirming that the phenotype we observed is more likely due to a disruption of assembly, rather than an upstream modulation of protein abundance (**Figure 4D**).

### SNED1 directly interacts with collagen I

The colocalization we observed between collagen I and SNED1 and the fact that SNED1 deposition in the ECM appears altered when collagen I is less abundant prompted us to test whether these two proteins interact. We performed biolayer interferometry and found that collagen I bound to Sned1 in a concentration-dependent manner (**Figure 4E**). To date, collagen I is the first experimentally identified partner binding directly to full-length SNED1. Interestingly, in performing these experiments, we found that covalently immobilized Sned1 did not bind soluble collagen I, suggesting that the conformation and flexibility of SNED1 likely influence its interaction with collagen I. In addition, as discussed above, the fact that soluble collagen I can directly interact with SNED1 does not warrant that these two proteins interact within the insoluble ECM scaffold. It would thus be interesting to probe the interaction between these two proteins *in situ* (Leverton et al., 2026).

Since the surface of collagen fibrils is decorated by other ECM components, such as small leucin-rich proteoglycans, it would also be interesting to determine the roles of proteoglycans and glycosaminoglycans in SNED1 assembly, since these molecules are known to modulate fibrillogenesis of ECM proteins (Hill et al., 2022; Kalamajski and Oldberg, 2010; Robinson et al., 2017). Last, whether this interaction is required to mediate the incorporation of SNED1 in the ECM in a fibrillar pattern remains to be determined. To test this, we will first need to characterize with greater details how SNED1 interacts with collagen I.

## CONCLUSIONS

The data presented in this short report are providing the first insights into the timeline of incorporation of SNED1 during the process of ECM build-up and into the mechanisms governing the fibrillar assembly of SNED1 in the ECM. While our results are demonstrating a key role for the presence of fibronectin and collagen I in the fibrillar assembly of SNED1, the precise molecular mechanisms by which they regulate SNED1 remain to be elucidated. It will also become important to determine whether the same mechanisms are at play *in vivo*, in particular in processes requiring SNED1 such as craniofacial morphogenesis during development and breast cancer metastasis.

## MATERIALS AND METHODS

### Plasmid constructs

The cDNA encoding full-length human SNED1 cloned into pCMV-XL5 was obtained from Origene (clone SC315884). The cloning of GFP-tagged SNED1 (hereafter referred to as SNED1-GFP) was previously described (Vallet et al., 2021). The construct was then shuttled into the bicistronic retroviral vector pMSCV-IRES-Puromycin between the BglII and EcoRI sites, as previously described (Vallet et al., 2021) and was used for all imaging experiments.

SNED^1-751^ was subcloned into p-Select-CGFP-Blasti (Invivogen) between the AgeI and NcoI restriction sites, and then shuttled into the bicistronic retroviral vector pMSCV-IRES-Puromycin between the BglII and EcoRI sites. The sequences of the primers used for subcloning are listed in **Supplemental Table S1**. All constructs were verified by Sanger sequencing at the UIC Genome Research Core facility.

### Cell culture

#### Cell maintenance

Stable expression SNED1^1-751^-GFP into immortalized mouse embryonic fibroblasts (iMEFs) isolated from *Sned1*^KO^ mice (*Sned1*^KO^ iMEFs) (Barqué et al., 2021) was performed as previously described for SNED1-GFP (Vallet et al., 2021). *Sned1*^KO^ iMEFs stably overexpressing GFP alone, SNED1-GFP, or SNED1^1-751^-GFP, and iMEFs isolated from *wild type* mice were cultured in Dulbecco’s Modified Eagle’s medium (DMEM; Corning, #10-017-CV) supplemented with 10% fetal bovine serum (FBS; Sigma, #F0926) and 2 mM glutamine (Corning, #25-005-CI), this medium will be further referred to as complete medium. All cell lines were maintained at 37 °C in a 5% CO_2_ humidified incubator.

#### Preparation of fibronectin-depleted FBS

Fibronectin was depleted from FBS using a gelatin Sepharose 4B affinity resin (Cytiva, #17095601) (Sabatier et al., 2009; Sechler et al., 1996). All steps were performed on ice or in a cold room using ice-cold reagents. Gelatin Sepharose resin was supplied in 20% ethanol and was washed with 1X PBS before packing. 100 mL of resin was packed into an Econo column using an Econo gradient pump (Bio-Rad) with a maximum flow rate of 6.0 mL/min. Once packed, the column was equilibrated with 5 column volumes of 1X PBS at a flow rate of 4.5 mL/min. Following equilibration, 300 mL of FBS was loaded onto the column at a flow rate of 4.0 mL/min using an Econo pump and looped for 200 minutes. The flow-through was sterilized using a 0.2µm filter. Pre- and post-chromatography samples were analyzed for the presence of fibronectin using immunoblotting.

#### RNA interference

iMEFs overexpressing SNED1-GFP were seeded on 12mm-diameter glass coverslips coated with 0.18 mg/mL Matrigel (Corning, #356234) inserted in non-cell-culture-treated 24-well plates. Cells were seeded at such a density to reach 60% confluency 2-4 hours post-seeding, in medium containing 10% of fibronectin-depleted FBS. 2-4 hours after seeding, cells were transfected using Lipofectamine 3000 (Invitrogen, #L3000008) with 5 pmol of control or fibronectin-targeting short interfering RNAs (siRNAs) per well (sequences provided in **Supplemental Table S3**). 48 hours after seeding, the culture medium was replaced with fresh medium containing 10% of fibronectin-depleted FBS. 72 hours post-seeding, cell cultures were decellularized and the resulting ECMs were fixed and stained to visualize ECM proteins as described below.

### Deoxycholate (DOC) solubility assay

The DOC solubility was performed as previously described (Pankov and Yamada, 2004; Wierzbicka-Patynowski et al., 2004). iMEFs overexpressing SNED1-GFP were seeded at 5.5 × 10^5^ in sterile cell culture-treated 6-well plates. Approximately 48 hours after seeding, 50 μg/mL ascorbic acid (Thermo Scientific, #352685000) was added to each well. After that, half of the medium was replaced with complete medium containing 100 μg/mL ascorbic acid every 48 hours. After a specified time in culture (3 days, 6 days, or 9 days), the conditioned medium (CM), containing proteins secreted by the cells, was collected and centrifuged for 3 minutes at 800 x *g* to remove any cellular debris, and the DOC solubility assay was performed on the cell culture layer composed of cells and their associated ECM. In brief, cells were washed with cold 1X PBS. 1 mL of DOC lysis buffer (5% deoxycholate, 3 M Tris-HCl pH 8.8, 500 mM EDTA), protease inhibitors with EDTA (Thermo Scientific, #A32953), and 167 μg/mL DNase I (Sigma, #AMPD1-1KT) was added to each well and incubated at 4°C for 1 hour. Samples were then collected using a sterile scraper and transferred to 1.5 mL tubes. Samples were sheared using a 26G needle to reduce viscosity then centrifuged for 15 minutes at 21,100 × *g* at 4 °C. The supernatant (DOC-soluble material) was transferred to a tube and stored at -80 °C. The pellet, containing DOC-insoluble proteins, was stored at -80 °C. Protein quantification of the DOC-soluble fraction was performed via BCA assay, according to the manufacturer’s instructions (Thermo Scientific, #23225). Samples were prepared for SDS-PAGE as previously described (Wierzbicka-Patynowski et al., 2004). In short, the concentrations of DOC-soluble protein fractions were normalized to the sample containing the lowest total protein concentration to ensure equal loading of protein in each lane (12-15 μg). Since the solubilization of DOC-insoluble protein fractions requires high detergent and reducing agent concentrations (6% SDS and 100mM DTT, respectively), their precise concentration cannot be measured. Thus, and as previously described (Wierzbicka-Patynowski et al., 2004), to normalize protein loading of DOC-insoluble protein fractions were solubilized in a volume of Laemmli buffer based on the total protein concentration of their respective DOC-soluble fractions.

In parallel, total protein extracts (TE) from cells seeded and treated under the same conditions were collected by lysing the cells in 3X Laemmli buffer (0.1875 M Tris-HCl pH 6.8, 6% SDS, 30% glycerol) containing 100 mM dithiothreitol. Samples were then sheared using a 26G needle to reduce viscosity and boiled for 10 minutes at 95 °C and stored at -80 °C.

### Immunoprecipitation of SNED1-GFP from conditioned medium

iMEFs overexpressing SNED1-GFP were seeded at 5.5 x 10^5^ in a sterile tissue-culture-treated 6-well plate. When cells had reached 100% confluency (∼48h post-seeding), the culture medium was replaced and supplemented with 50 μg/mL ascorbic acid. The conditioned medium (CM) was collected 4 days post-seeding and subjected to immunoprecipitation using protein A/G agarose beads (Thermo Scientific, #20421) and either IgG isotype as a negative control (Invitrogen, #PI31903) or an anti-GFP antibody (Sigma, #G6539; **Supplemental Table S2**). A second negative control, omitting antibodies, was also included. Samples were incubated on a rotating wheel at 4 °C, overnight. Samples were then centrifuged at 8,200 x *g* at 4 °C for 30 seconds to pellet the beads and immunoprecipitated proteins. Beads were washed three times in cold 1X PBS. Following the final centrifugation, beads were dried and immunoprecipitated proteins were resuspended in 3X Laemmli buffer containing 100 mM DTT. Samples were boiled for 10 minutes at 95 °C and then analyzed by SDS-PAGE and immunoblotting (*see below*).

### SDS-PAGE and immunoblotting

Protein samples were separated by SDS-PAGE at constant current (20 mA for the stacking gel, 25 mA for the separating gel). Proteins were then transferred to nitrocellulose membranes at 100V (constant voltage) for 3 hours at 4 °C. Membranes were stained with Ponceau stain for 10 minutes and imaged to confirm the transfer of proteins. Membranes were then blocked in 5% non-fat milk in PBS containing 0.1% Tween-20 (PBST) for 60 minutes at room temperature under constant shaking. Membranes were incubated with primary antibodies (**Supplemental Table S2**) in 5% non-fat milk in PBST overnight with constant shaking at 4 °C. Membranes were washed three times with PBST and then incubated with horseradish peroxidase (HRP)-conjugated secondary antibodies (**Supplemental Table S2**) in 5% non-fat milk in PBST for 60 minutes with constant shaking at room temperature. Membranes were again washed three times with PBST. The presence of proteins of interest was detected by chemiluminescence, using the Pierce ECL Western blotting substrate (Thermo Scientific #32109). Imaging was performed using a ChemiDoc MP imaging system (Bio-Rad).

### Generation of cell-derived ECMs

#### Cell seeding

Cell-derived ECMs from iMEFs overexpressing GFP alone or SNED1-GFP were prepared under aseptic conditions following a previously established protocol (Harris et al., 2018) with minor modifications. Briefly, sterile glass coverslips were inserted in non-cell-culture-treated sterile plastic plates and coated with autoclaved and sterile-filtered 0.2% gelatin in 1X Dulbecco’s phosphate buffered saline with calcium and magnesium (hereafter referred to as D-PBS^++^) (Cytiva, #SH30264.01) for 1 hour at 37 °C. The gelatin solution was removed, and coverslips were washed with 1X D-PBS^++^. The gelatin coating was crosslinked using 1% glutaraldehyde (Electron Microscope Sciences, #16220) in PBS for 30 minutes at room temperature. The glutaraldehyde solution was removed, and coverslips were washed 3x5 minutes with 1X PBS. To quench any remaining glutaraldehyde, 1 M ethanolamine (Thermo Scientific, #451762500) in MilliQ water was added to each well and incubated for 30 minutes at room temperature. The ethanolamine solution was removed, and coverslips were washed 3x5 minutes with 1X PBS. After the final wash, complete medium was added to each well, and coverslips were stored at 37 °C until cell seeding. 2.0 x 10^5^ cells per 18 mm-diameter coverslip or 1.0 x 10^5^ cells per 12 mm-diameter coverslip were seeded. When cells reached 100% confluency (approximately 48 hours after seeding), the medium was replaced and supplemented with 50 μg/mL ascorbic acid. After that, half of the medium was replaced with complete medium containing 100 μg/mL ascorbic acid every 48 hours. To assess the effect of withholding ascorbic acid on SNED1 assembly (-AA condition), cells were instead treated with D-PBS^++^ instead of ascorbic acid at the same frequency as the ascorbic acid treatment.

#### Decellularization

After a specified time in culture (3, 6, or 9 days), cell layers were decellularized. Cells were first washed twice in 1X PBS, then washed three times using pre-warmed wash buffer 1 (100 mM Na_2_HPO_4_, 2mM MgCl_2_, 2 mM EGTA, pH 9.60). Cells were then incubated with pre-warmed lysis buffer (8 mM Na_2_HPO_4_, 3% IGEPAL CA-360, pH 9.60) for 15 minutes at 37 °C. After the lysis buffer was added, the plate was tilted to aid in cell lysis. Tilting was repeated every 5 minutes, for a total of three times. The lysis buffer was replaced with a second and then a third incubation with fresh, pre-warmed lysis buffer, for 40 minutes at 37 °C. Samples were then incubated with pre-warmed wash buffer 2 (10 mM Na_2_HPO_4_, 300 mM KCl, pH 7.50) with 0.5% deoxycholate (Thermo Scientific, #J6228822) for 3 minutes and then washed three times with wash buffer 2, followed by four washes in deionized water at room temperature. 1X PBS was added to coverslips, and the samples were kept protected from light in aluminum foil at 4 °C until further processing. After each step, coverslips were visualized using a light microscope to assess cell removal.

### Immunofluorescence staining of cell-derived ECMs

Decellularized cell-derived ECMs were fixed with 4% paraformaldehyde (PFA; Fisher Scientific, #50980487) for 15 minutes at room temperature. PFA was quenched using 50 mM NH_4_Cl in PBS for 15 minutes at room temperature. Following fixation, samples were rinsed three times with 1X D-PBS^++^ and stained immediately or stored in 1X PBS at 4 °C for future use.

#### Staining of decellularized ECMs

Decellularized and fixed ECMs were stored for 96 hours at 4 °C in 1X PBS. Samples were blocked with 1% bovine serum albumin (BSA) in PBS containing 0.1% Triton X-100 at room temperature for 1 hour to prevent non-specific binding. Samples were then incubated with primary antibodies (**see Supplemental Table S2**) in 1X PBS containing 0.05% Triton X-100 overnight at 4 °C. Coverslips were washed three times with 1X D-PBS^++^ containing 0.1% Triton X-100 and then incubated with secondary antibodies in 1X PBS + 0.05% Triton X-100 for 1 hour at room temperature. Coverslips were washed three times with 1X D-PBS^++^ + 0.1% Triton X-100, then mounted on glass microscope slides using Fluoromount G. The slides were stored at 4 °C overnight prior to imaging.

### Imaging

#### Image acquisition

Samples were imaged using a Zeiss Confocal LSM 880 microscope.

#### Image analysis

##### ECM thickness analysis

Z-stacks obtained via confocal microscopy were analyzed to measure the thickness of the layers made of different proteins in decellularized ECMs. The bottom-most (beginning) and top-most (end) slices, where in-focus signal was observed, were recorded. Thickness was calculated by taking the total thickness of the Z-stack divided by the total number of slices to obtain the thickness of each individual slice. The beginning slice number was subtracted from the end slice number, and the resulting value was multiplied by the slice thickness (0.33-0.35 μm) to obtain the protein layer thickness value. These values were calculated for 3 to 5 fields per coverslip, for each protein.

##### 3D reconstruction

The volume of each Z-stack was visualized using Fiji ImageJ. Volume Viewer plugin was used to analyze stacks, with the Z-aspect set to 10.0 in Max Projection mode with Tricubic smooth interpolation. XY, XZ, and XY snapshots were obtained for each field.

##### Colocalization analysis

Colocalization analysis was performed on Z-stacks using Zeiss ZEN. Thresholds were set using negative controls, and analyses between channels for each Z-slice were recorded using Microsoft Excel. Line graphs using the Manders correlation coefficient were then plotted to visualize the changes across the specified portion of each Z-stack.

##### Area fraction density analysis

ECM density/area fraction was measured using Fiji ImageJ. 8-bit orthogonal projection (maximum intensity) images were imported and brightness and contrast were adjusted manually. Area fraction was selected in the analyze > set measurements menu to obtain a value of staining density for 3 to 5 independent fields per coverslip (*see respective Figure legends*).

### Biolayer interferometry

The binding of recombinant His-tagged full-length mouse Sned1 (R23-K1403) expressed in Chinese hamster ovary cells (Bio-Techne, #9335-SN) to collagen I was investigated by biolayer interferometry (BLI) in an Octet RED96 system (Sartorius), as previously described . Human acid-soluble collagen I from skin (Sigma, #C5483) was solubilized at 0.5 or 1 mg/ml in 0.5 M acetic acid and used for the assay in a 10X kinetics buffer containing PBS pH 7.4, 0.02% Tween 20 and 0.1% BSA (Sartorius) plus 0.6 M sucrose, 20 mM imidazole and 1% BSA (BIS buffer). Recombinant mouse Sned1) was dissolved in kinetics buffer 10X (Sartorius) at 125 µg/ml, diluted to 5 µg/ml in PBS, pH 7.4, and captured for 5 minutes on Ni-NTA sensors (Sartorius) via its C-terminal His tag. Control sensors were prepared by omitting Sned1 and replacing it with PBS. After the capture step, the sensors were washed with BIS buffer for 3 minutes. SNED1-coated and control sensors were dipped for 5 minutes in a solution of collagen I diluted in BIS buffer at 26-150 µg/ml or 87.5-500 nM. Non-specific binding to the sensor surface (*i.e.*, the signal recorded on control sensors) was subtracted from the signals recorded on SNED1-coated sensors to get specific binding. The binding levels were measured for 5 seconds before the dipping end, and the complexes formed on the sensor surface were allowed to dissociate for 10 minutes in BIS buffer. BLI experiments were performed three independent times.

### Statistical analysis

All experiments were performed with at least three biological replicates, unless otherwise indicated. Data is represented as individual experimental values, mean ± standard deviation or standard error of the mean, as indicated in the respective figure legends. Dot plots were generated using GraphPad Prism. As indicated in each figure legend, Welch and Brown-Forsythe one-way ANOVA with Dunnett’s T3 correction for multiple comparisons or unpaired t-tests with Welch’s correction were performed using GraphPad Prism to assess any statistical differences across groups and/or conditions.

## ACKNOWLEDGEMENTS

We would like to thank all the members of the Naba lab for insightful discussions, James Considine and Dr. Amanpreet Kaur Bains for their technical help, and Dr. Christophe Quétard (Sartorius) for his valuable advice on the BLI experiments.

## DATA AVAILABILITY

All relevant data can be found within the article and its supplemental information. Research materials are available upon request to Dr. Naba.

---

**Supplemental Figure S1.**
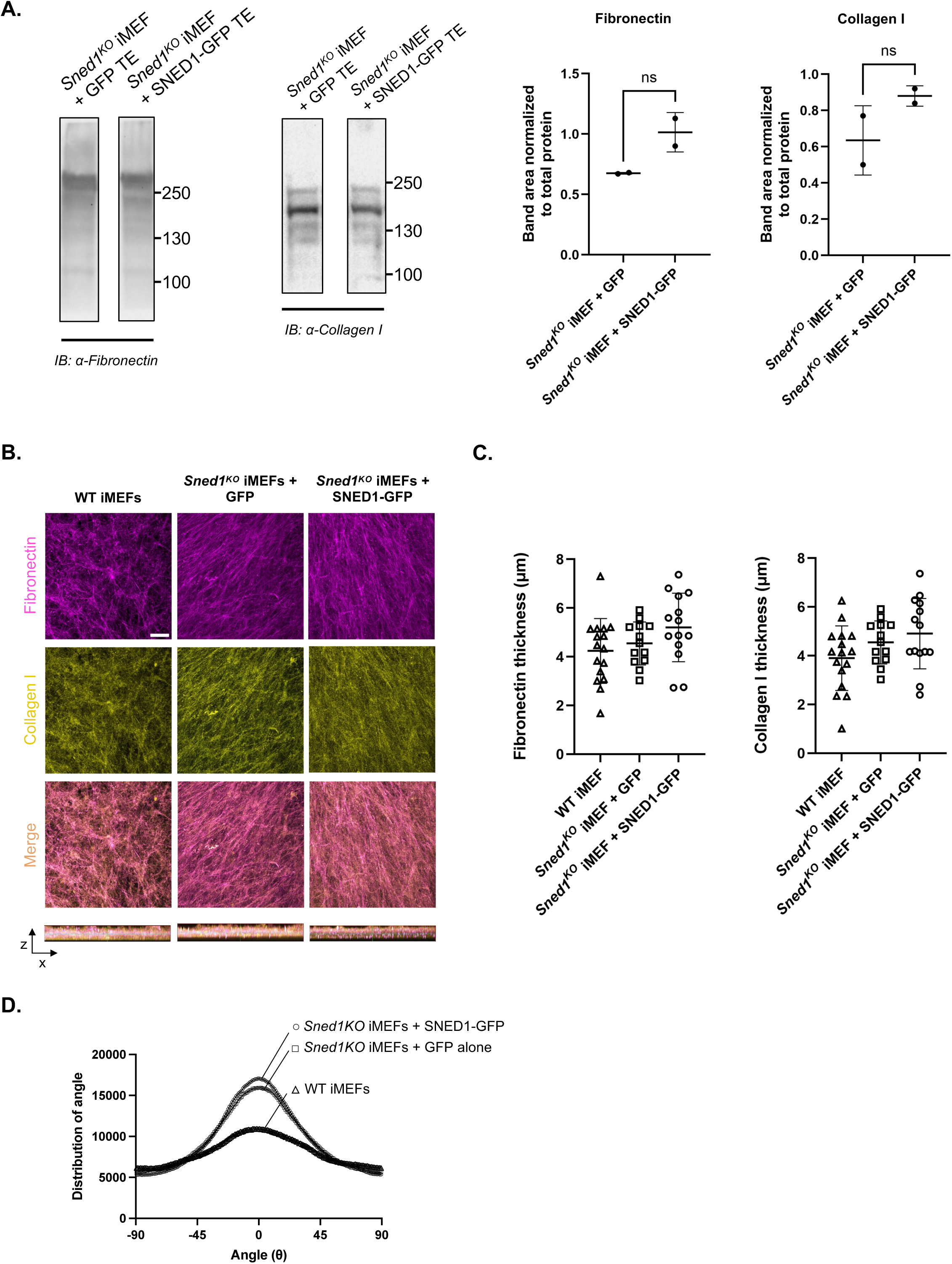
Overexpression of SNED1-GFP does not alter fibronectin or collagen I assembly in the ECM; *related to Figure 1*. A. *Left panels:* Immunoblots using an anti-fibronectin or an anti-collagen I antibody show the presence of the respective proteins in total protein extracts collected from *Sned1^KO^*iMEFs expressing GFP alone or SNED1-GFP. ***Right panels:*** Bar charts represent the quantification of immunoblot signal intensity, where the area of the bands corresponding to fibronectin or collagen I in total protein extracts of *Sned1^KO^* iMEFs expressing GFP alone or SNED1-GFP were normalized to total protein content determined by Ponceau staining (N=2). Unpaired t-test with Welch’s correction did not show statistical significance (ns). **B.** Orthogonal maximum projections (XY, *top panels*; XZ, *lower panels*) show fibronectin (magenta) and collagen I (yellow) in the ECM produced by wild-type (WT; *left panels*) iMEFs, *Sned1^KO^* iMEFs expressing GFP (*middle panels*), and *Sned1^KO^* iMEFs overexpressing SNED1-GFP (*right panels*) and decellularized 9 days post-seeding. Scale bar: 20 µm. Images are representative of at least three biological replicates with at least 4 fields imaged per replicate and per timepoint. **C.** Dot plots compare the thickness of the fibronectin (*left panel*) and collagen I (*right panel*) signal (µm) in decellularized ECMs produced by wild-type (WT) iMEFs, *Sned1^KO^* iMEFs expressing GFP, and *Sned1^KO^*iMEFs overexpressing SNED1-GFP, and decellularized 9 days post-seeding. Individual experimental values from three independent biological replicates, with at least 4 fields per replicate and timepoint, are represented together with the mean value (black bar) ± standard deviation. Welch and Brown-Forsythe one-way ANOVA with Dunnett’s T3 correction for multiple comparisons shows no statistically significant differences between the groups. **D.** Histograms depict fibronectin fiber alignment in ECMs produced by WT iMEFs (triangles), *Sned1^KO^* iMEFs overexpressing GFP (squares), or *Sned1^KO^* iMEFs overexpressing SNED1-GFP (circles), decellularized 9 days post-seeding. Data are represented as the mean of three biological replicates, with at least 4 fields per replicate.

**Supplemental Figure S2.**
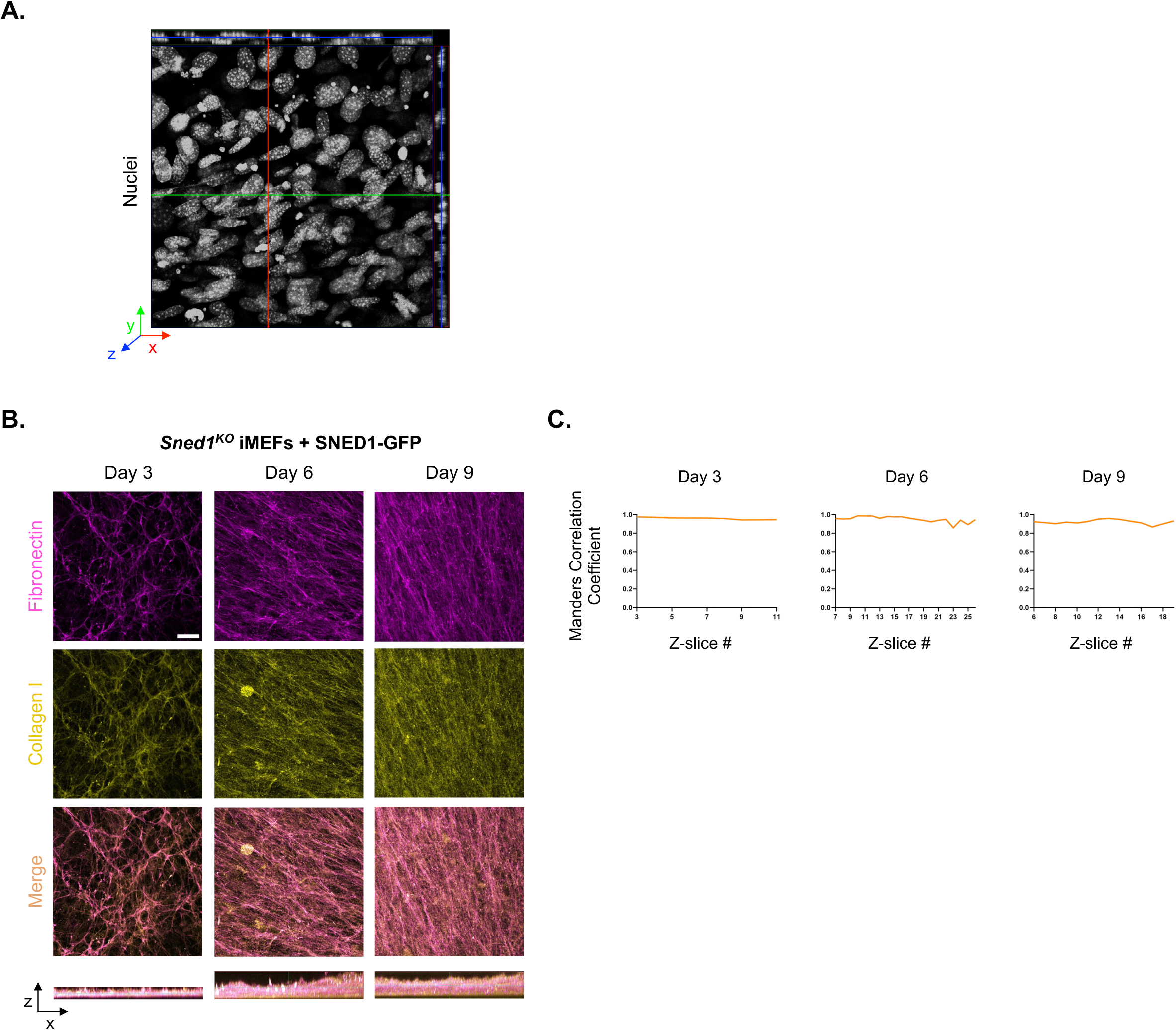
Expression of SNED1-GFP does not alter the colocalization of fibronectin or collagen I; *related to Figure 2*. **A.** Orthogonal projections of XY, XZ and YZ show the distribution of nuclei of *Sned1^KO^*iMEFs overexpressing SNED1-GFP in culture for 9 days and prior to decellularization. The image is representative of at least three biological replicates. The X-axis is represented in red, the Y-axis in green, and the Z-axis in blue. **B.** Orthogonal maximum projections (XY, *top panels*; XZ, *lower panels*) show the merge of fibronectin (magenta; see single-stain panel in **Figure 2A**) and collagen I (yellow; see single-stain panel in **Figure 2E**) in the ECM produced by *Sned1^KO^*iMEFs overexpressing SNED1-GFP and decellularized 3, 6, or 9 days post-seeding. Scale bar: 20 µm. Images are representative of at least three biological replicates with at least 4 fields imaged per replicate and timepoint. **C.** Line graphs show, for each timepoint, the Manders overlap coefficient between the fibronectin and collagen I signals in the ECM produced by *Sned1^KO^* iMEFs overexpressing SNED1-GFP for each Z-slice of a representative imaging field.

**Supplemental Figure S3.**
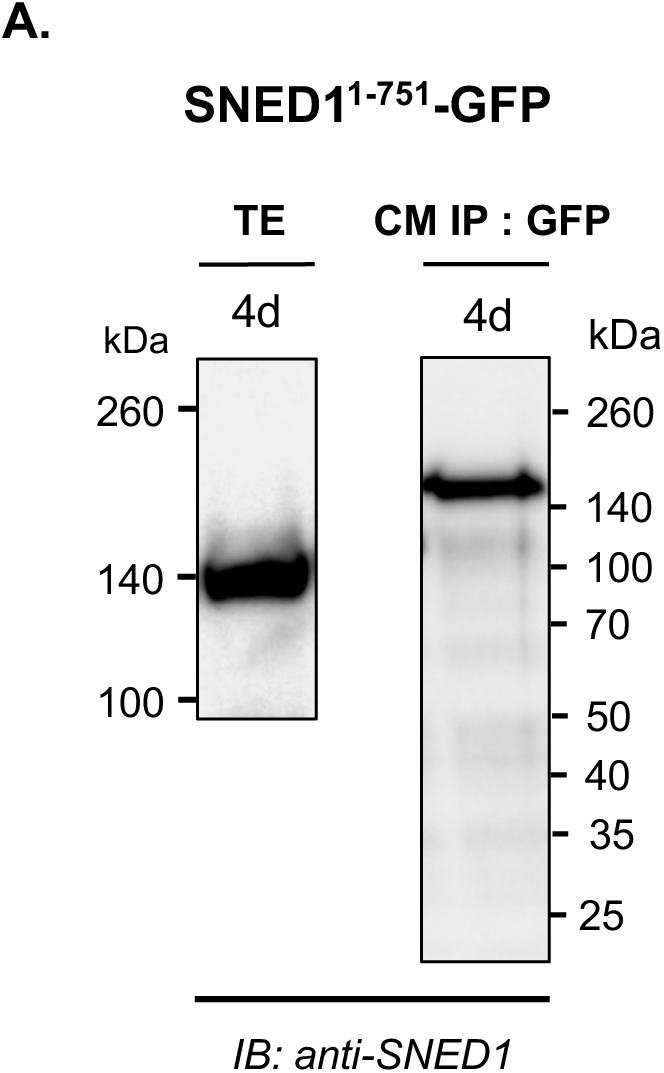
Expression of SNED1^1-751^-GFP in immortalized *Sned1^KO^* iMEFs; *related to Figure 3E*. A. Immunoblots show the presence of SNED1^1-751^-GFP in the total protein extract (*left panel*) and conditioned medium (*right panel*) of *Sned1^KO^* iMEFs overexpressing SNED1^1-751^-GFP.

**Supplemental Figure S4.**
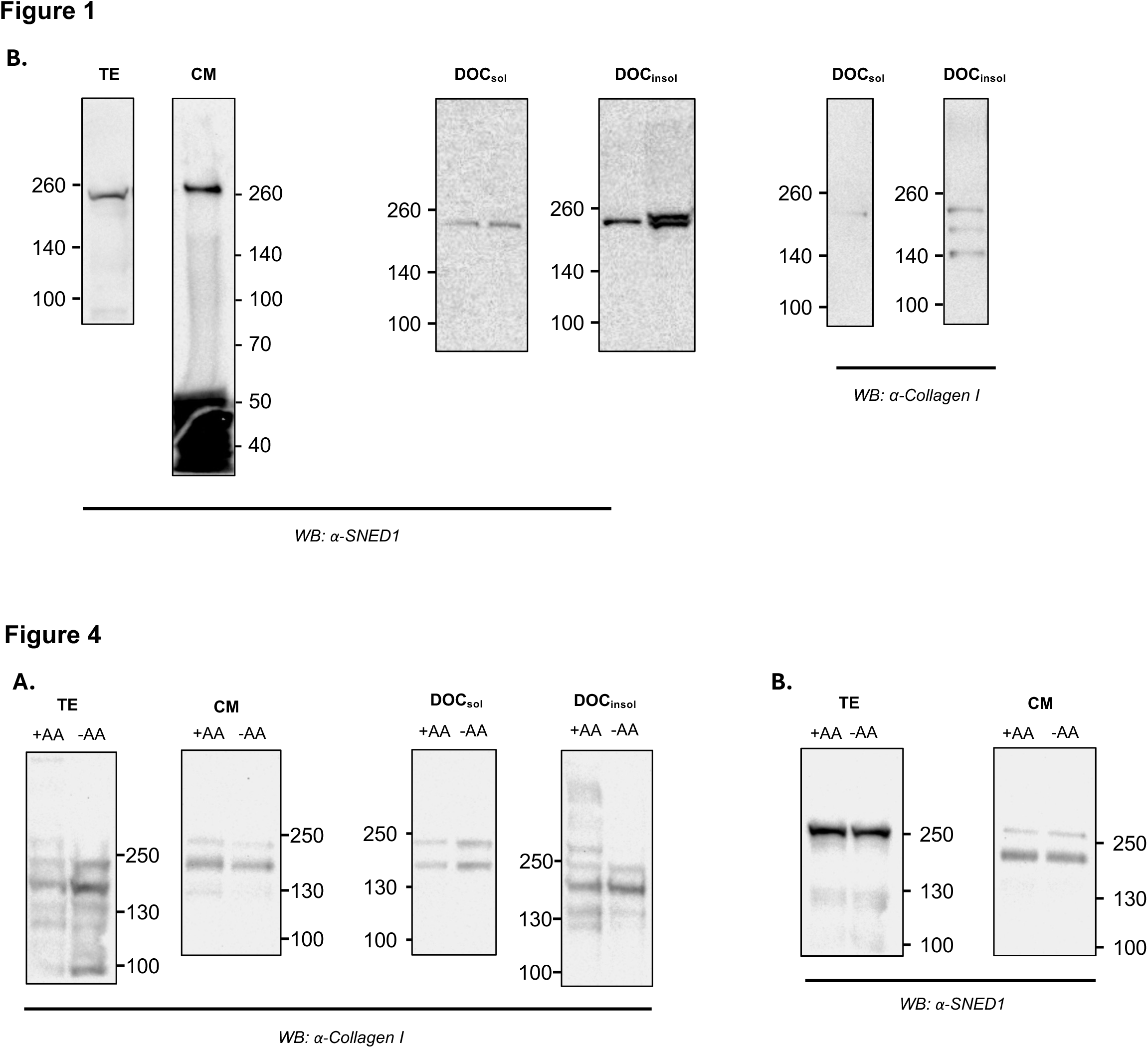
Immunoblot transparency. Uncropped immunoblots corresponding to all panels presented in the main figures are provided.

**Supplemental Table S1:**
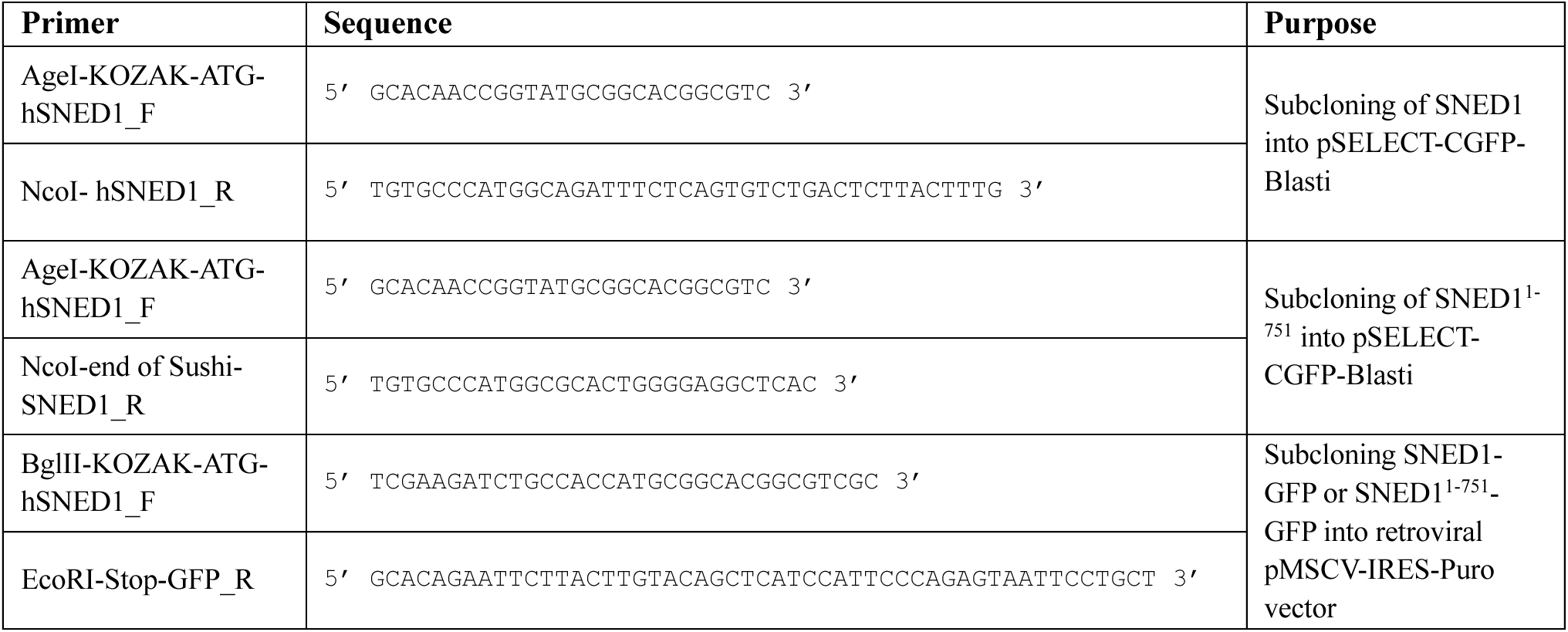
List of primers.

**Supplemental Table S2:**
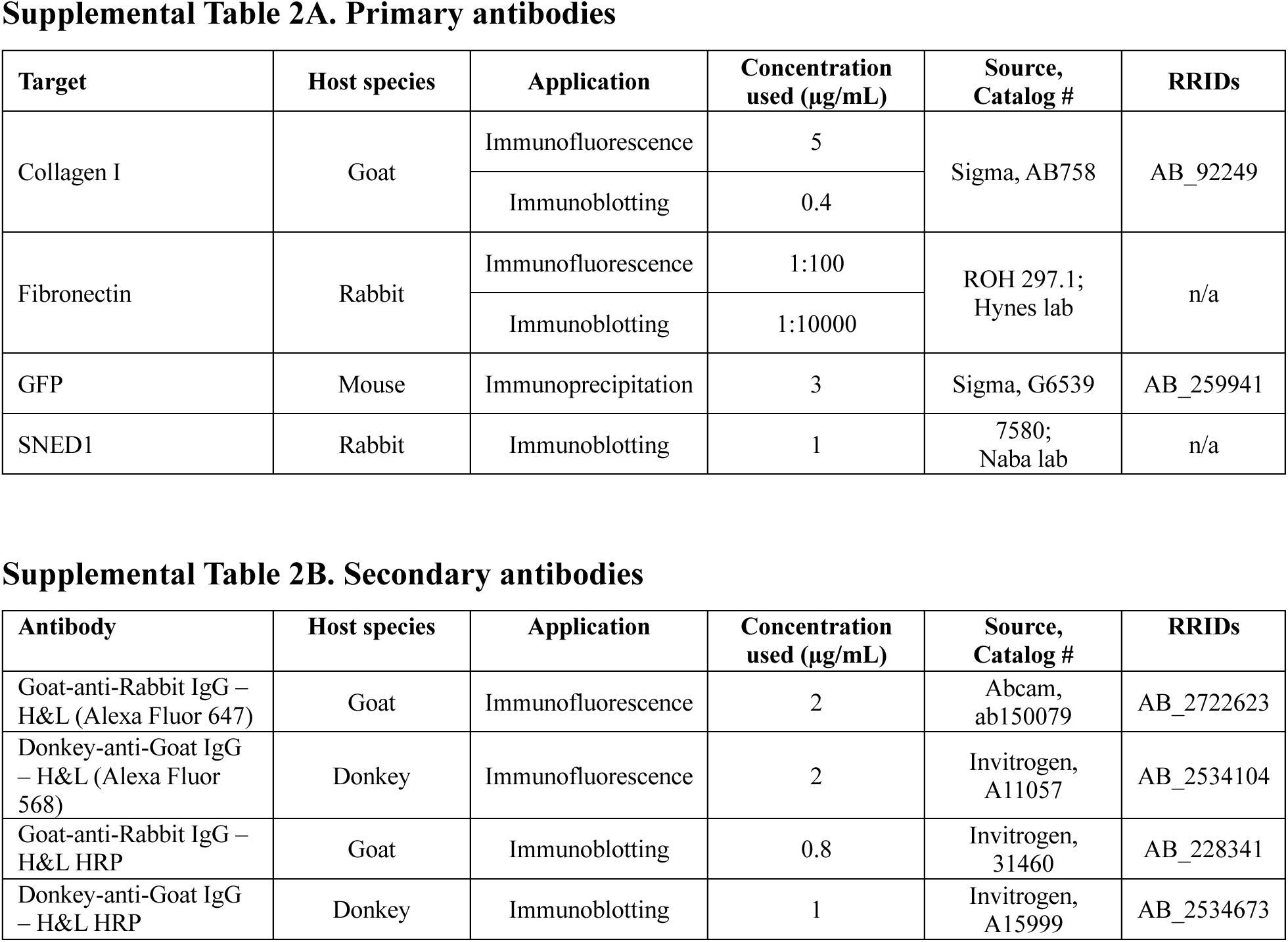
List of antibodies used for immunostaining and immunoblotting.

**Supplemental Table S3:**
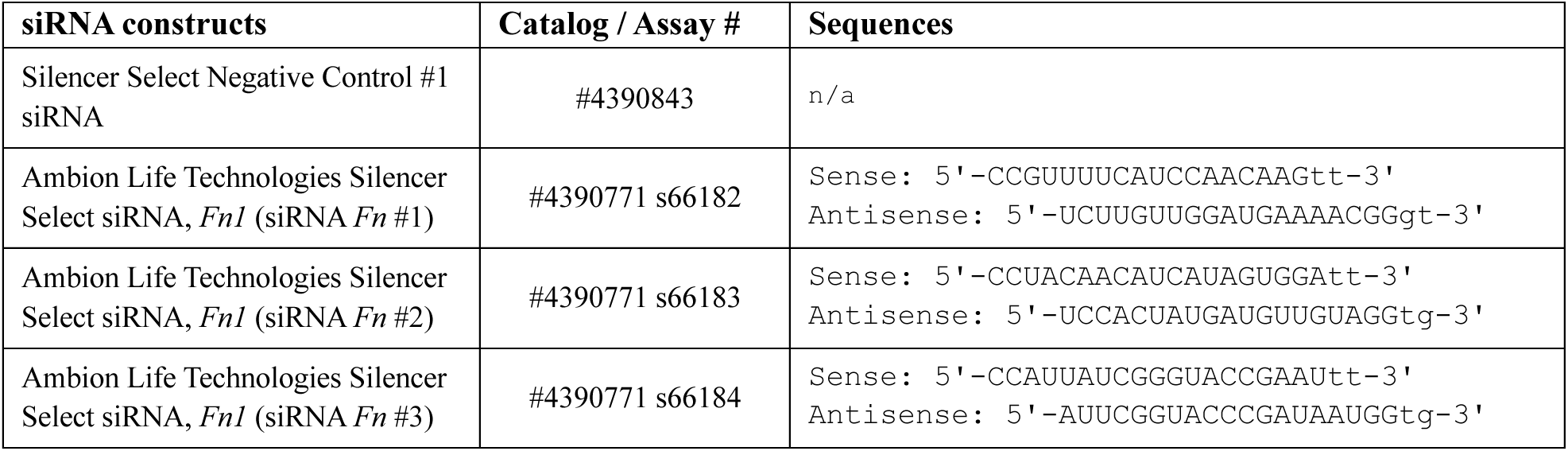
siRNA sequences.

